# A Portable Software Library that Implements the Gillespie Direct Method

**DOI:** 10.1101/181446

**Authors:** Herbert M Sauro

## Abstract

A common approach to simulating chemical reaction systems is to formulate the problem as a set of ordinary differential equation and solve them using standard numerical procedures. Such a formulation works well when the number of molecules involved in the simulation is large and assumes a deterministic formulation for the underling chemical kinetics. However for systems such as gene regulatory networks where the number of molecules of a specific species can be of the order of 10s of molecules, the deterministic approach fails to capture important dynamical features. At low molecular numbers stochastic events dominate the dynamics. As a result much research has been devoted to developing numerical algorithms for computing the stochastic trajectories. One of the most common approaches is based on Gillespie’s stochastic simulation algorithm, for example the direct method. Due to its simplicity the direct method has been implemented many times in software. However, unusually, there does not appear to be a C based reusable software library. This article describes the development of such a library. All source code is licensed under the liberal Apache 2.0 open source license and is available at Github: https://github.com/sys-bio/libStochastic

## 1 Introduction

A common approach to simulating chemical reaction systems is to formulate the problem as a set of ordinary differential equation and solve them using standard numerical procedures. Such a formulation works well when the number of molecules involved in the simulation is large and assumes a deterministic formulation for the underling chemical kinetics. However for systems such as gene regulatory networks where the number of molecules of a specific species can be of the order of 10s of molecules, the deterministic approach fails to capture important dynamical features. At low molecular numbers stochastic events dominate the dynamics. As a result much research has been devoted to developing numerical algorithms for computing the stochastic trajectories. One of the most common approaches is based on Gillespie’s stochastic simulation algorithm, for example the direct method. Due to its simplicity the direct method has been implemented many times in software. However, unusually, there does not appear to be a C based reusable software library. This article describes the development of such a library.

## 2 Algorithm

The Gillespie direct method algorithm itself has been described many times in the literature. A good source of information on this is the paper by McCollum et al. [5] who describe a variety of approaches, including comparing the relative performance of each. The table below summaries the direct method algorithm (modified from McCollum):

~~~
n = number of species in the model
Initialize vector S of species amounts
Initialize vector R of rate constants
currentTime = 0.0
while currentTime endTime do
begin
totalPropensity = 0.0
for i = 0 to n
begin
propensity[i] = calculatePropensities (S, R)
totalPropensity = totalPropensity + propensity[i]
end
dt = -ln (rand())/totalPropensity
selector = totalPropensity * rand()
for i = 1 to n
selector = selector -propensity[i]
if selector = 0 then
begin
selectedReaction = i
break
end
end
// Update Reactants and Products of selected reaction
for all reactants of selectedReaction do
reactants[selectedReaction] = reactants[selectedReaction] -reactantStoichiometry
for all products of selectedReaction do
products[selectedReaction] = products[selectedReaction] + productStoichiometry
output_Results()
currentTime = currentTime + dt
end
~~~

The algorithm first computes all the propensity functions (reaction rates). It then draws two uniformly distributed random numbers. It uses one of the random numbers to select which reaction will ‘fire’ and the second random number to determine when it will ‘fire’ in the future as a Δ*t*. The reactants and products for the selected reaction are updated, taking into account the corresponding stoichiometries. The process is then repeated.

A formal as well as an intuitive understanding of the direct method can be found in the enzyme kinetics text book by Sauro [7].

## 3 Currently Available Software

There is a range of software available for computing stochastic trajectories. However none of the current software can be easily reused. For the purpose of this article, reusable software is considered open source software (using a non-restrictive license) packaged into a platform and language independent software library. By platform independent we mean that the source code must not include any platform or operating system specific code. For example any external libraries should be readily available on all platforms and any file operations should be made through platform independent means. By language independent we mean that the library can be used from any other programming language and not just the language that the library is written in. This eliminates, for example computer languages such as Java or C#.

A reusable library would allow stochastic simulations to be carried out in a variety of other applications, including scripting languages such as Python or Julia or other programming languages such as C++, Object Pascal, Go, Rust etc.

The direct method has been implemented in most software languages including unexpected applications such as Excel. An examination of the literature and repositories such as Github or sourceforge reveals many implementations. Some implementations are embedded in large applications and therefore cannot be easily removed for reuse. Some are written in languages that are difficult to embed in other languages, for example, C#, Java, Python, Matlab or Julia. There is at least one library, StockKit [6], that is written in C++ however the code is licensed under the strict GPL license which makes the software almost impossible to distribute with non-GPL code. StockKit however supports a great variety of other approaches for example tau-leaping, and automatic parallelism.

In this article a new library called libStochastic is described that is written in standard C (C99) https://en.wikipedia.org/wiki/C99.

## 4 libStochastic Library

libStochastic is a new library written in standard C (C99). The library implements the direct method and supports a liberal open source license (Apache 2.0) and a simple C based API. The use of C allows the library to be used by virtually any other computer language. As an example, a simple Python interface is described later in the article but the library could equally be interfaced to other systems such as R, Julia Java, Object Pascal etc. SWIG (http://www.swig.org/) can be used to wrap the library into over twenty-three other languages.

The libStochastic library consists of nine files. Most of these contain code to support simple data structures such as dynamic arrays and matrices. One file, gl_Stochastic contains the implementation of the direct method. The direct method requires access to a random number generator. It is possible to use the standard random number generator supplied with the C runtime libraries. However the more robust Mersenne-Twister [4] is provided instead. This is an implementation of MT19937 [3], meaning it has a period of 2^19937^ *-* 1. The version included with the library is one developed by Christian Stigen Larsen http://csl.name. The original code can be found at https://github.com/cslarsen/mersenne-twister.

### Using the Library

The core library implements the standard direct method for generating stochastic trajectories. The library supports the reaction types shown in Table 1. At present the library does not support arbitrary rate laws though this could be added at a later date.

To initialize a simulation run an instance of the direct method algorithm must first be created. This is accomplished using the method gl_createStochastic(). The method returns a handle of type gl_STOCHASTIC OBJ that should be used in all subsequent calls. Once a simulation has been completed, the memory associated with the handle should be released using the method: gl_freeStochastic(). For example:

gl_STOCHASTIC_OBJ gl = gl_createStochastic();

// …Specify model, run simulation and collect data…

gl_freeStochastic (gl);

**Table 1:**
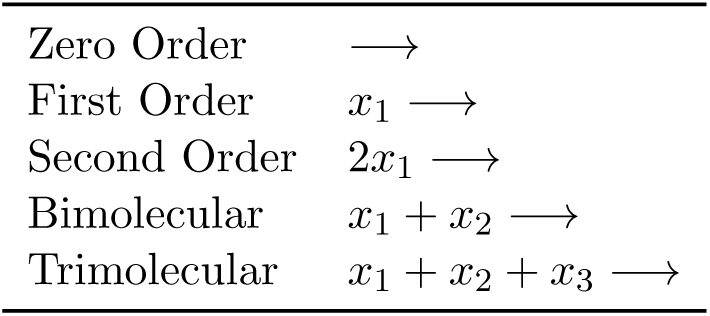
Allowed reaction types. Note that any number of products are permitted in a given reaction.

Three simulation methods are provided, gl_execute, gl_executeOnGrid and gl_moveOneStep. gl_execute will generate a single trajectory given a start and end time. The last point generated will be the penultimate point before reaching the specified end time. The method returns a matrix that contains the simulation data. The first column of the matrix will always be time, the remaining columns represent the amounts of the floating species.

For example: m = gl_execute (gl, 0, 5.0)

Due to the nature of the algorithm, the times that are reported in the first column of the matrix are irregular. Often however it is desirable to generate simulation data on a regular grid. For this purpose the library supplies the alternative method, gl_executeOnGrid. This method takes an additional argument which represents the number of points, The time distance between a point is computed using *dt* = (*timeEnd -timeStart*)*/*(*numberOf Points -* 1). For example if timeEnd and timeStart are set to 0 and 5 respectively and the number of points set to 6, then output will be generated at points, 0, 1, 2, 4, and 5. Generating simulation runs at fixed intervals can be useful when the average of many runs is needed or when fewer data points are required.

For finer control, the library also supplies gl_moveOneStep. A call to this routine will execute a single time step. The method returns the new time. This call can be useful for developing more specialized data collection. For example if rate constants or amounts need to be changed mid flight due to an external signal. Two methods, gl_setAmount and gl_getAmount are provided to set or get the amounts of a specific species in the model. Likewise, gl_setRateConstant can be used to set the value of a rate constant in a specific reaction.

Two other methods that are useful when using the execute methods. These are gl_reset and gl_setInitialAmount. When the method gl_addSpecies is called during model construction, one of the arguments is the initial amount of the species which is stored is a specific initalAmounts array. The actual amounts that are changed during the simulation are stored in a separate runtime array. When gl_execute or gl_executeOnGrid is called the simulation starts from the *current state* of the model as stored in the runtime array. In order to start a subsequent simulation using the initial conditions, the gl_reset method can be called which loads the runtime array with the contents of the initial amounts array. The advantage of this approach is that repeated calls to gl_execute or gl_executeOnGrid will continue the simulation where the previous call left off. At any time a simulation needs to be called again using the original initial amounts, gl_reset can be called. Finally, gl_setInitialAmount allows one to change the initial amounts.

**Table 2:** List of methods used to add reactions to a model

|  |  |
| --- | --- |
| Add a Zero Order Reaction | `gl_addZeroOrderReaction` |
| Add a First Order Reaction | `gl_addFirstOrderReaction` |
| Add a Second Order Reaction | `gl_addSecondOrderReaction` |
| Add a UniUni Reaction | `gl_addUniUniReaction` |
| Add a Bimolecular Reaction | `gl_addBimolecularReaction` |
| Add a Trimolecular Reaction | `gl_addTrimolecularReaction` |
| Add a product | `gl_addProduct` |

**Specifying a Model** A variety of methods are provided to specify a model. The library does not support modeling standards such as SBML directly but this could be easily added by way of the Python interface. To specify a model the first operation to do is add the species in the model. To illustrate how a model is built consider the following reaction model:

~~~
xo -> x1; 0.1
x1 -> x2; 0.6
x2 -> x1; 0.3
x2 ->; 0.15
~~~

In this model we assume that the amount of *x*_*o*_ is **fixed** and does not change during the simulation. This is to illustrate the use of fixed boundary species. The two middle reactions form a cycle and the last reaction is used to illustrate a first order reaction that drains into unspecified sink. The numbers on the right-hand side are the rate constants (sometimes called the reaction hazards). The model contains three species which can be added to the model using the method gl_addSpecies:

~~~
gl_STOCHASTIC_OBJ gl = gl_createStochastic();
xo = gl_addSpecies (gl, ’xo’, false);
x1 = gl_addSpecies (gl, ’x1’, true);
x2 = gl_addSpecies (gl, ’x2’, true);
~~~

The third argument indicates whether the species has a fixed level or is variable (floating). Note that C99 supports the words, true and false, unlike ANSI C. Of importance is that that gl_addSpecies returns a integer index that can be used to refer to the species when adding reactions to the model. It is therefore useful to be able to store these values are we generate them.

Once the variables have been added to the model the reactions can be included. There are six methods for adding reactions. These are listed in Table 2

The gl_addproduct method should be used to add products to a reaction and their corresponding stoichiometry. Any number of products can be added to a reaction. For example it can be used to specify reactions such as *x*_1_ *→ x*_2_, *x*_1_ *→ x*_2_ + *x*_3_, *x*_1_ *→* 2*x*_2_ + 10*x*_2_ + 3*x*_3_, or even a reaction such as *x*_1_ *→ x*_1_ which effectively models a fixed species.

If products are unspecified, then reactions of the type *x →* can be described i.e where the reaction drains into an anonymous sink.

Two short-cut methods (addUniUniReaction and addBioMolecularReaction) are provided if it is known that the reaction only produces a single product with a stoichiometry of one. These reactions include *x*_1_ *→ x*_2_, 2*x*_1_ *→ x*_2_ and *x*_1_ + *x*_2_ *→ x*_3_.

The method gl_addUniUniReaction will add a reaction of the form *x*_1_ *→ x*_2_, for example: gl_addUniReaction (g, x1, x2, 0.5)

The method gl_addBiUniReaction will add either 2*x*_1_ *→ x*_2_ or *x*_1_ + *x*_2_ *→ x*_3_, depending on the arguments, for example:

~~~
gl_addBiUniReaction (x1, x2, x3, 0.4) Adds x1 + x2 -> x3
gl_addBiUniReaction (x1, x1, x2, 0.4) Adds 2 x1 -> x2
~~~

Returning to the example described earlier, the reactions can now be specified. Note that in the method calls, the species indexes returned from gl_addSpecies are used. All the reaction methods other than gl_addProduct return a reaction index. This index should be used when adding products to the reaction. The last reaction added does not include a gl_addPRoduct call because this reaction is of the form *x*_2_ *→*.

~~~
gl_REACTION *rxa;
rxa = gl_addFirstOrderReaction (gl, xo, 0.1);
gl_addProduct (rxa, x1, 1);
rxa = gl_addFirstOrderReaction (gl, x1, 0.6);
gl_addProduct (rxa, x2, 1);
rxa = gl_addFirstOrderReaction (gl, x2, 0.3);
gl_addProduct (rxa, x1, 1);
gl_addFirstOrderReaction (gl, x2, 0.15);
~~~

Note: In all cases, incorrectly specifying the species or reaction index will result in an error condition. In general methods will return a −1 in the event of such an error.

### Data Handling

The gl_execute and gl_executeOnGrid methods return the results in the form of a custom matrix. Three methods are provided to access the data in the matrix and include gl_getMatrixValue, gl getMatrixRows and gl_getMatrixCols. The last two methods retrieve the number of rows and columns in the matrix respectively. This information combined with gl_getMatrixValue can be used to access all the data in the matrix. Once the data has been extracted, the memory used by the matrix should be released by calling gl_freeMatrix.

### Interfacing to Python

To use the library it is necessary to call the library from a host application. Because the library exposes a pure C API, the library can be called from almost any other programming language including of course another C or C++ application. However it is becoming increasingly common to use a scripting language such as Python to use external libraries. The advantage is that performance critical parts of the application can be written in a compiled language such as C whereas the more often tedious data handling and display can be handled by more easy to use languages such as Python. Many scripting language now support an extensive set of libraries as well as comprehensive plotting capabilities. The added advantage of using Python is that it is extremely easy to learn and is an ideal vehicle for novice users to gain access to more sophisticated algorithms.

There are a variety of approaches to linking a C library to Python, the three common approaches include ctypes, SWIG and the use of specific Python embedding code in the library. The later is of no interest because it requires Python specific changes to the C library which would render the library Python only. The two other methods are the use of ctypes and SWIG. SWIG is a general purpose tool that allows a library with a C compatible API (or C++) to be automatically ‘wrapped’ and made available to a wide range of other languages including Python. For long term development SWIG is an excellent solution but it requires some experience to use including a compilation stage. In this article ctypes will be used because it doesn’t require any compilation and if necessary it can be used directly from the Python console. The main drawback of ctypes is that one has to be careful when specifying the types as it is possible to crash the Python runtime.

### Handling External Events

It is sometimes the case that during a simulation an external event occurs at a specific time. For example a parameter or species amount is changed. Events can be implemented by stitching together multiple simulations. For example, assume that a reaction parameter k1 must be doubled at a specific time, say at 25 time units and then the simulation continued. This can be accomplished with the following calls to the library:

~~~
// C Code:
// set up model k1 = 2.5
….
// When creating the reaction, save the reaction index
// so that later on we can change the rate constant
reactionIndex = …..
// Run first part of the simulation
// Args: library handle, time start, time end, number of points
m1 = gl_executeToGrid (gl, 0, 25, 51)
gl_setRateConstant (gl, reactionIndex, 2*k2)
// Run the second half of the simulation
m2 = gl_executeToGrid (gl, 25, 50, 51)
~~~

Note the time start argument in the second simulation call has been adjusted to 25.

The current Python bindings were developed as a quick way to test the library and as a result lack some important error checking. A more long term solution would be to create formal SWIG libraries for a number of languages, include Python, R, Scilab and Octave.

## 5 Testing

Evans et al. [1] developed a stochastic simulation test suite that include 39 test models in all. Some of these are focused on testing SBML compliance but a number of the tests focus on the stochastic algorithm itself. The problem with testing stochastic simulators is that each simulation run is unique. It is not possible therefore to just compare a single run an the expected outcome. Instead it is necessary to compare the distribution of outcomes after running a large number of simulations and averaging across all trajectories. The Evans test suite includes the expected means and standard deviations for each of the test models which allows us to compare the expected outcomes to the simulation run. The Evans tests also provide a means to make a statistical assessment of how well the expected outcomes match the simulated data. The authors recommend that *at least* 10,000 trajectories are evaluated for each test. All tests should be carried out using the executeOnGrid call so that individual trajectories can be readily matched up across each time point. To conserved memory, the tests implement a running average and standard deviation. There are standard methods for computing running statistical measures and are designed to avoid numerical instability. Calculating a cumulative mean is straight forward and is given by:

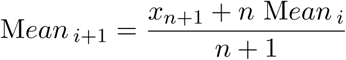

where *x*_*i*_ is the *i*^*th*^ data point and *n* the number points so far processed.

For computing the cumulative variance, the recommended method is to use an algorithm developed by Welford and popularized in Donald E. Knuth’s book the The Art of Computer Programming, volume 2: Seminumerical Algorithms, 3rd edn (1998). This is given below:

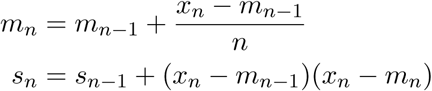

where *m*_1_ = *x*_1_ and *s*_1_ = 0. The sample variance is given by *s/*(*n -* 1). One additional advantage of this approach is that the mean, *m* is computed at the same time. In practice the mean and variance is computed at every point along a trajectory by computing across all points at a given time. These calculations can be be very conveniently and succinctly represented using Python by employing the numpy library which can carry out arithmetic operations on vectors:

~~~
# Python :
import numpy as np
# Set up the empty matrices first, first column is time (which we ignore),
# but the gl_execute methods generates a time column so we need to allocate
# space for it. The remaining columns are the individual species in the model
M = np.zeros(shape=(numberOfPoints, numberOfVariables+1))
S = np.zeros(shape=(numberOfPoints, numberOfVariables+1))
# Generate the first trajectory
# Mp will be used to represent the previous (k-1) trajectory
Mp = gl.executeOnGrid(0.0, timeEnd, numberOfPoints)
# Generate many trajectories
for i in range (sizeOfEnsemble):
# simulate again with same initial conditions
gl.reset()
data = gl.executeOnGrid(0.0, timeEnd, numberOfPoints)
# Divide by i+2 because i starts at zero
# and we’ve already had one processed
np.copyto (M, Mp + (data -Mp)/(i+2))
np.copyto (S, S + (data -Mp)*(data -M))
# Make the current data the previous
np.copyto (Mp, M)
# Finally compute the variance
variance = variance/(sizeOfEnsemble-1)
~~~

The full Python code is available on Github https://github.com/sys-bio/libStochastic. It may not be the most elegant code but it works. As suggested before, not all the Evans’ tests are applicable. For example 001-02 and 001-08 and others are SBML dependent tests, that is they ensure that the SBML model is being interpreted correctly. Since we don’t read the models from SBML, such tests are not applicable. Another set of tests which don’t currently use are due to limitation of libStochastic and relate to use of arbitrary propensity functions which are not currently supported. These include tests such as 001-013 and 001-014 as well as others. Related to this is test 001-019 which requires an assignment rule which would include arbitrary math. Again libStochastic does not currently support evaluating arbitrary math. The final limitation is that libStochastic does not support volumes other than unity. This eliminates test 001-018. This leaves fourteen tests out of a total of thirty-nine. Of the fourteen, libStoachastic passes every test.

### Memory Leak/Corruption Tests

Tests for memory leaks were carried out using Microsoft’s CRT debug heap manager by setting the CrtSetDbgFlag. This will identify memory leaks, heap corruption, and buffer overrun errors. One leak and one potential heap corruption bug was identified using this method. Both bugs were eliminated in the final release.

## 6 Examples

In this section four examples are shown that illustrate the use of the library. All runs were carried out via the Python bindings and matplotlib was used to plot the data.

## Equilibration Model

The equilibration model is simply:

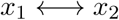

The python code for this model is give below:

~~~
# Python:
from stBindings import *
import pylab
gl = Stochastic ()
x1 = gl.addSpecies (’X1’, 80, True)
x2 = gl.addSpecies (’X2’, 0, True)
rxa1 = gl.addFirstOrderReaction (x1, 0.1)
gl.addProduct (rxa1, x2, 1)
rxa2 = gl.addFirstOrderReaction (x2, 0.3)
gl.addProduct (rxa2, x1, 1)
m = gl.execute (0, 100)
pylab.plot (m[:,0], m[:,1], color=’r’)
pylab.plot (m[:,0], m[:,2], color=’b’)
pylab.savefig (’equilibration.pdf’)
gl.free()
~~~

The script saves the plot to a pdf file which makes it easy to import to other applications (for example LaTeX). Other formats such as png can also be saved. Figure 1 shows the resulting plot. pylab is imported to provide access to matplotlib.

**Figure 1:**
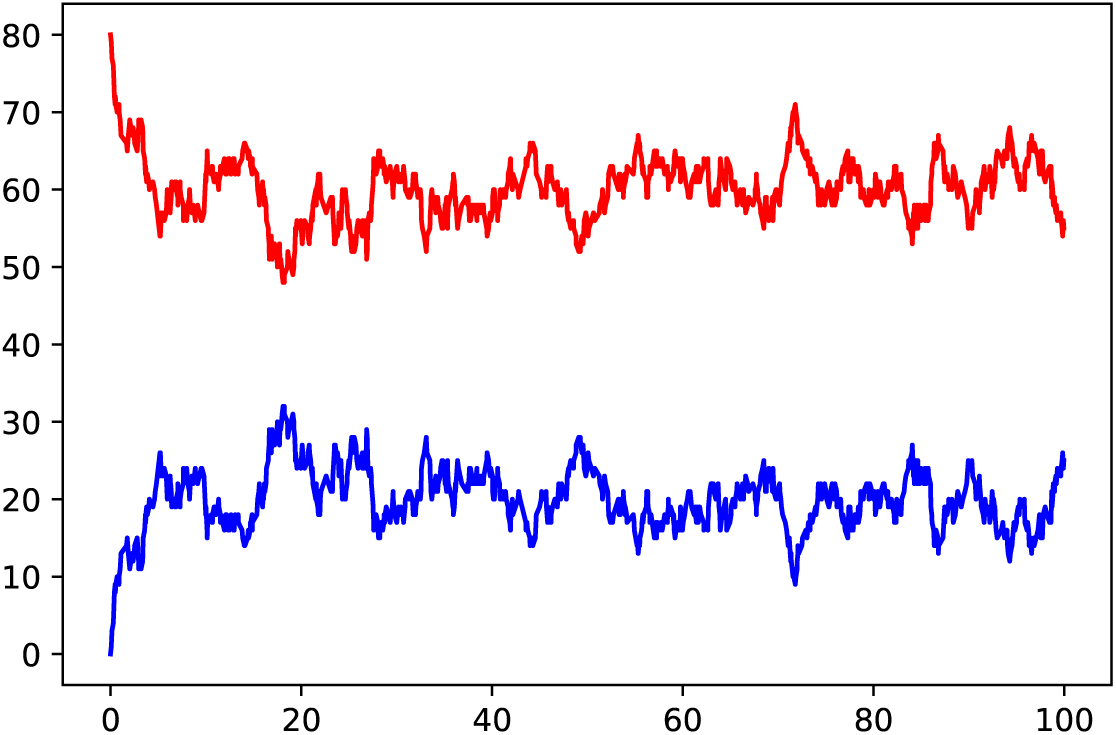
Equilibration example showing a single trajectory set.

The second example is a model of three consecutive but irreversible reactions: *x*_1_ *→ x*_2_ *→ x*_3_ *→ x*_4_. This time a ensemble of 60 models is generated and plotted simultaneously. A transparency of 20 percent is applied to each plot. In addition a *x* title is added to the plot (Figure 2).

~~~
# Python:
from stBindings import
* import pylab
gl = Stochastic ()
x1 = gl.addSpecies (’X1’, 50, True)
x2 = gl.addSpecies (’X2’, 0, True)
x3 = gl.addSpecies (’X3’, 0, True)
x4 = gl.addSpecies (’X4’, 0, True)
rxa1 = gl.addFirstOrderReaction (x1, 0.1)
gl.addProduct (rxa1, x2, 1)
rxa2 = gl.addFirstOrderReaction (x2, 0.3)
gl.addProduct (rxa2, x3, 1)
rxa3 = gl.addFirstOrderReaction (x3, 0.2)
gl.addProduct (rxa3, x4, 1)
for i in range (60):
gl.reset()
m = gl.execute (0, 70)
pylab.plot (m[:,0], m[:,1], color=’r’, alpha=0.2)
pylab.plot (m[:,0], m[:,2], color=’b’, alpha=0.2)
pylab.plot (m[:,0], m[:,4], color=’c’, alpha=0.2)
pylab.xlabel (’Time’)
pylab.savefig (’ThreeReactions.pdf’)
gl.free()
~~~

In the third example, the previous model is modified so that the source species, *x*_1_ and sink species *x*_4_ are fixed boundary species. In this situation the system with approach a steady state, albeit with fluctuations around the steady state, see Figure 3.

~~~
from stBindings import *
import pylab
gl = Stochastic ()
x1 = gl.addSpecies (’X1’, 80, False)
x2 = gl.addSpecies (’X2’, 0, True)
x3 = gl.addSpecies (’X3’, 0, True)
x4 = gl.addSpecies (’X4’, 0, False)
rxa1 = gl.addFirstOrderReaction (x1, 0.1)
gl.addProduct (rxa1, x2, 1)
~~~

**Figure 2:**
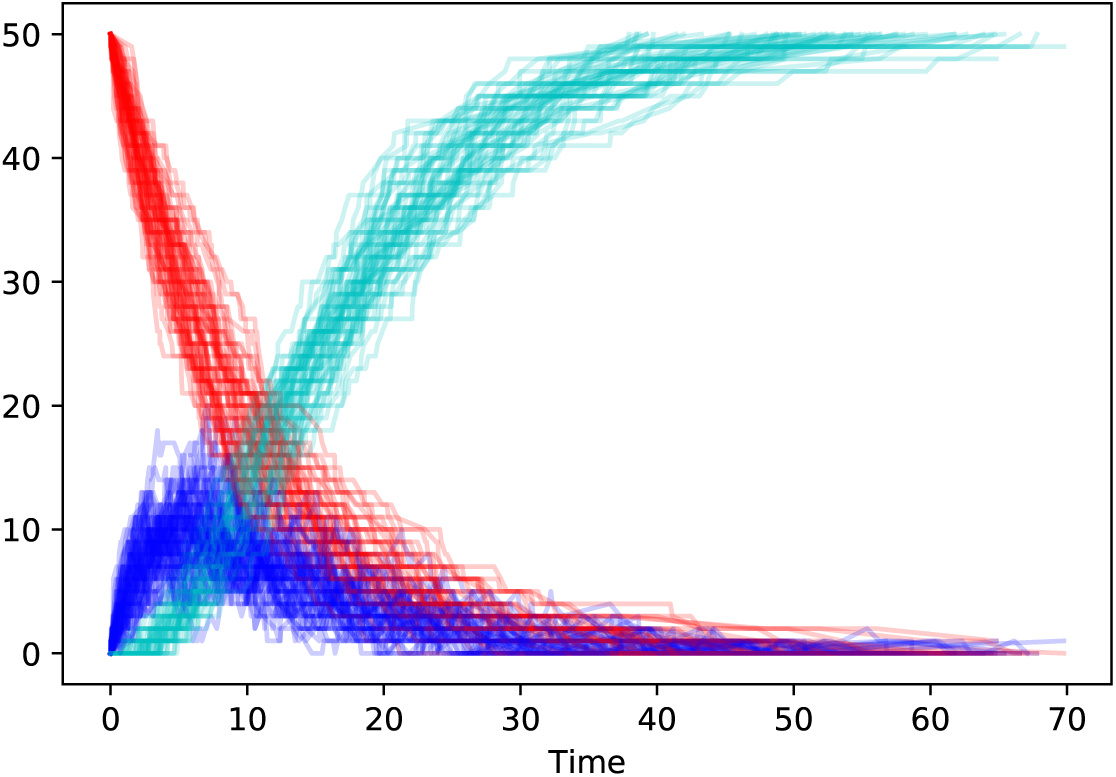
Three consecutive irreversible reactions where 60 independent plots are superimposed on each other.

~~~
rxa2 = gl.addFirstOrderReaction (x2, 0.65)
gl.addProduct (rxa2, x3, 1)
rxa3 = gl.addFirstOrderReaction (x3, 0.2)
gl.addProduct (rxa3, x4, 1)
pylab.figure(figsize=(8,5))
pylab.xlim((0, 70))
for i in range (60):
gl.reset()
m = gl.execute (0, 70)
pylab.plot (m[:,0], m[:,1], color=’r’, alpha=0.2)
pylab.plot (m[:,0], m[:,2], color=’b’, alpha=0.2)
pylab.xlabel (’Time’)
pylab.savefig (’ThreeReactionsSteadyState.pdf’)
gl.free()
~~~

In the last example the method executeOnGrid is used to simulate an event at a given time point. In this case the rate constant for the first reaction is doubled when time = 50.

~~~
from stBindings import *
import pylab
import numpy as np
gl = Stochastic ()
~~~

**Figure 3:**
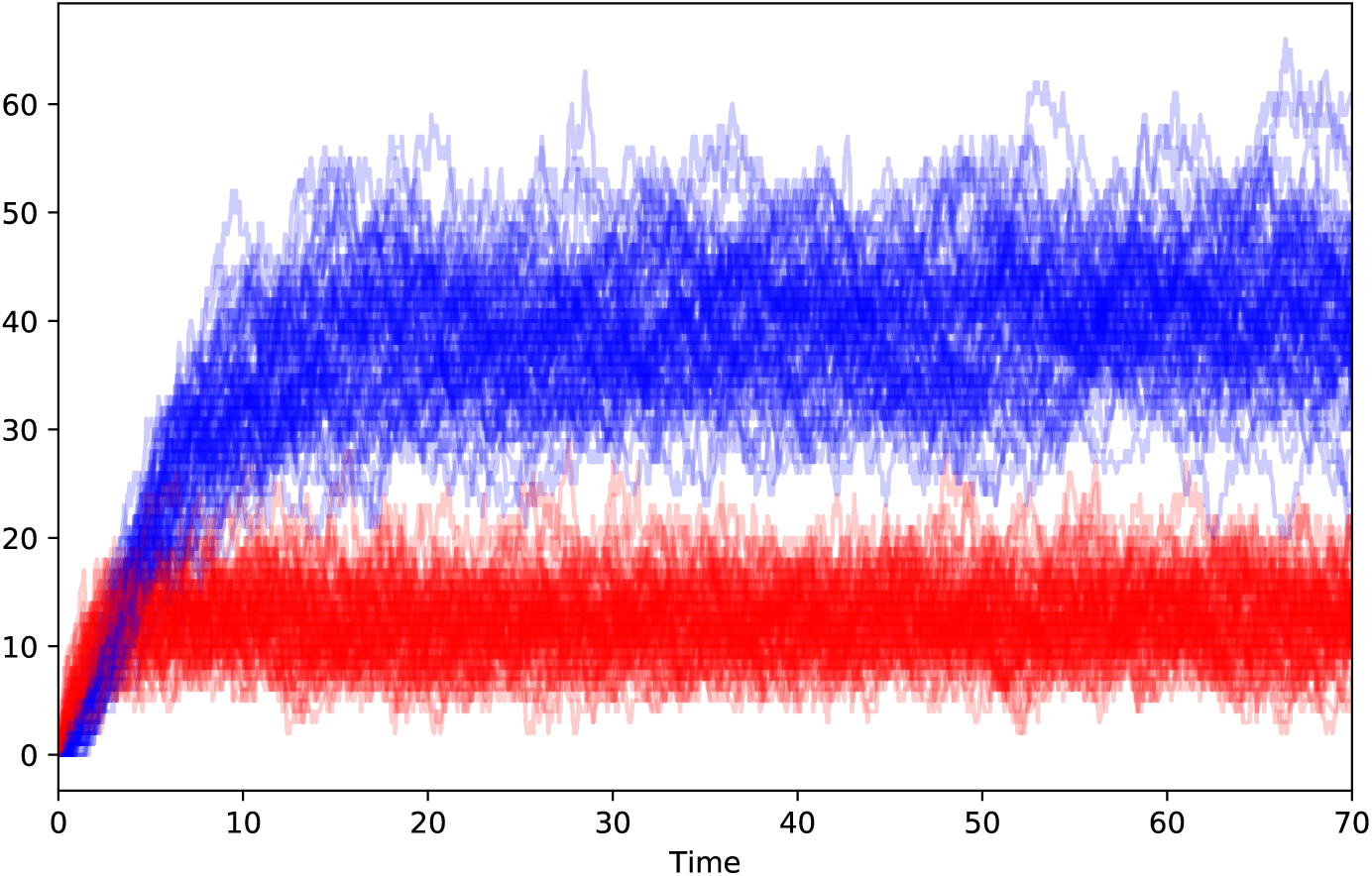
Three consecutive irreversible reactions where the source and sink are fixed boundary species, allowing the system to reach a state state. Due to the stochastic nature of the model the system fluctuates around the steady state. The plot shows 60 superimposed and independent trajectories.

~~~
x1 = gl.addSpecies (’X1’, 250, False)
x2 = gl.addSpecies (’X2’, 0, True)
x3 = gl.addSpecies (’X3’, 0, True)
x4 = gl.addSpecies (’X4’, 0, False)
rxa1 = gl.addFirstOrderReaction (x1, 0.1)
gl.addProduct (rxa1, x2, 1)
rxa2 = gl.addFirstOrderReaction (x2, 0.65)
gl.addProduct (rxa2, x3, 1)
rxa3 = gl.addFirstOrderReaction (x3, 0.2)
gl.addProduct (rxa3, x4, 1)
gl.setSeedUsingTime()
pylab.figure(figsize=(8,5))
pylab.xlim((0, 200))
m1 = gl.executeOnGrid (0, 100, 100)
gl.setRateConstant (rxa1, 0.2)
m2 = gl.executeOnGrid (100, 200, 100)
m3 = np.vstack ((m1, m2))
pylab.plot (m3[:,0], m3[:,1], color=’r’, alpha=1)
pylab.plot (m3[:,0], m3[:,2], color=’b’, alpha=1)
pylab.xlabel (’Time’)
pylab.savefig (’EventExample.pdf’)
gl.free()
~~~

The trick to implement an event is to carry out two simulations, the first goes from time 0 time 100 and the second from time 100 to time 200. When the first simulation is done, the rate constant for the first reaction is double. The second simulation is then started. Once complete, the data sets from the two simulations are stitched together to form one seamless run using vstack from the numpy library. A legend is also added to the plot. See Figure 4.

**Figure 4:**
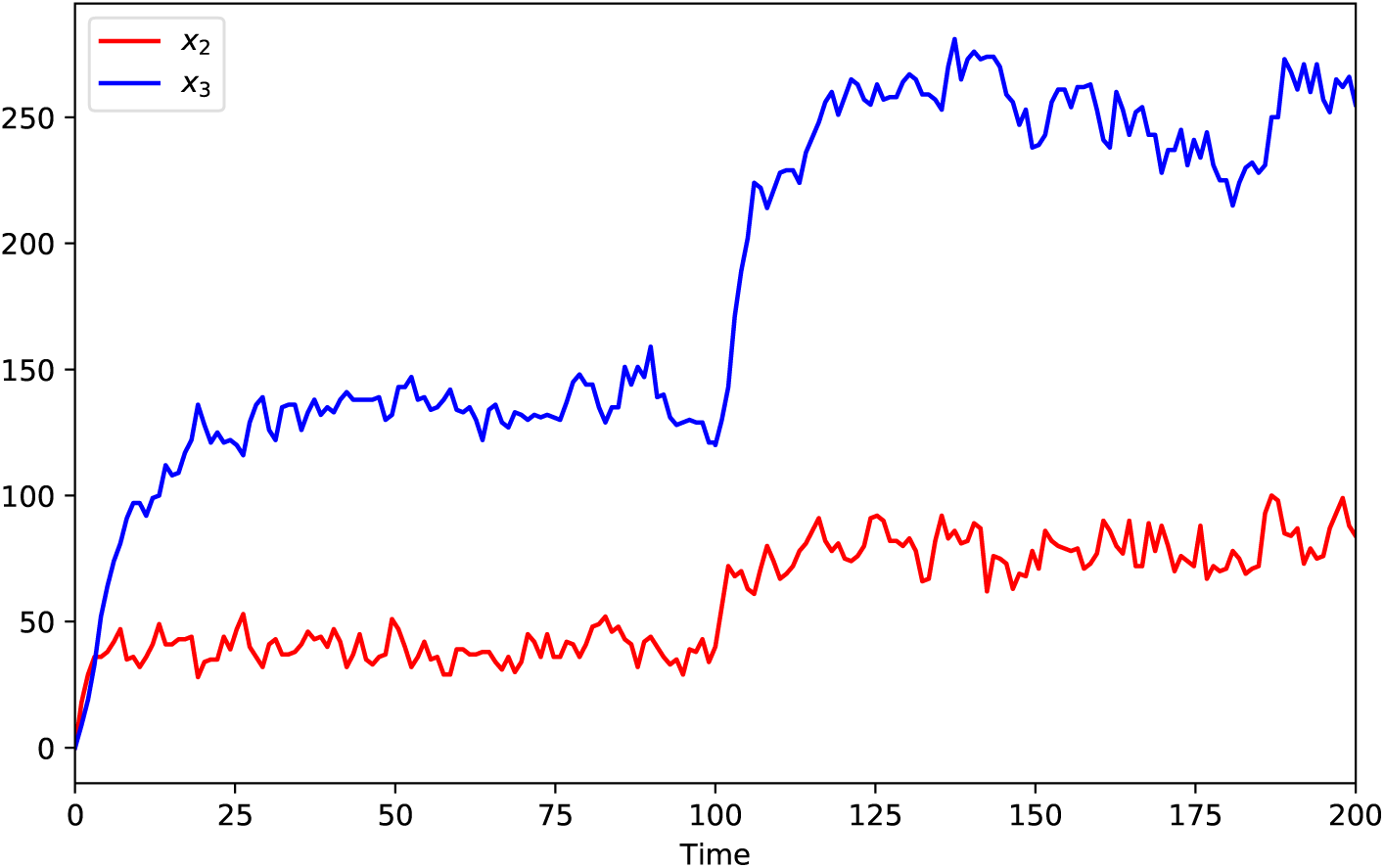
Three consecutive irreversible reactions where the source and sink species, *x*_1_ and *x*_4_ are fixed. A a time = 100, the rate constant for the first reaction is doubled and the simulation continued.

In addition to rate constants it is also possible to change amounts of boundary or floating species by using the gl_setAmount. The method requires the index of the species when it was originally created with gl_addSpecies.

## 7 Discussion

In this article a new C based portable chemical reaction stochastic simulator library is described. The library implements the standard direct method as described previously by Gillespie [2]. It defines an easy to use API that supports both standard trajectory generation using the direct method as well as generating results on a fixed time grid. This makes it easy to run ensembles and generate means and standard deviations. In addition it makes it easy to add events, to change parameters or species amounts to a simulation by stitching together multiple runs with events in between.

In addition, the moveOneStep method is exposed that allow users much more control over their simulations.

The software has been specifically written so that it is both platform and language agnostic. C99 is chosen as the implementation language which ensures that the library can be compiled and used by any other software host on any of the common operating systems. An example is shown where the library is called from Python. This binding is used to write the Python code to test the numerical correctness of the implementation.

The use of C99 in place of the older ANSI C allowed the use of C99 standard bool to represent the Boolean type. Other than that, the code is standard ANSI C.

There are some features missing which could be added at a later date. The most notable omission is the ability to set arbitrary math expression for propensities. One could argue that the direct method can not generate reliable simulation results for arbitrary expression, for example Michaelis-Menten or Hill equations. However, most modern implementations appear to permit users to specify almost any propensity expression.

## 8 Availability

All code and test models are available on Github and licensed under The Apache 2.0 open source license to allow maximum flexibility. At present the code has only been tested on windows using Visual Studio 2017 but given that the library uses standard C and doesn’t use any third-party code other than the Mersenne-Twister, there should be no issues in compiling the code on Linux or the Mac.The structure of the library is simple, one main file libStochastic.cpp and a number of supporting files that provide facilities for dynamic structures

## 9 Appendix

Sample C code that simulates a simple two step pathway that include a fixed source and sink and a single intermediate species, x.

~~~
#include <stdio.h>
#include “gl_Stochastic.h”
void main (int argc, char* argv[]) {
double timeStart = 0.0
double timeEnd = 50.0;
printf(“Starting Gillespie simulation…\n”);
gl_STOCHASTIC_OBJ *gl = gl_createStochastic();
int x = gl_addSpecies (gl, “x", 0, true);
int source = gl_addSpecies (gl, “source", 0, false);
int sink = gl_addSpecies (gl, “sink", 0, false);
int rxa = gl_addZeroOrderReaction (gl, 10.0);
gl_addProduct (gl, rxa, x, 1);
rxa = gl_addFirstOrderReaction (gl, x, 0.1);
gl_addProduct (gl, rxa, sink, 1);
gl_MATRIX *m = gl_execute (gl, timeStart, timeEnd);
for (int i = 0; i < m->nRows; i++) {
for (int j = 0; j < m->nCols; j++) {
printf(“%f “, m->m[i][j]);
}
printf(“\n”);
}
gl_freeMatrix(m);
gl_freeStochastic (gl);
printf (“Finished. “);
getchar();
}
~~~

## Acknowledgements

This work is supported by the National Science Foundation, under grants ABI 1355909, and MCB-1158573.

